# High fat diet promotes obesity by driving miRNA dependent neurogenesis and integration of orexigenic neurons in the feeding circuitry

**DOI:** 10.1101/2023.12.22.573029

**Authors:** Balakumar Srinivasan, Sarbani Samaddar, Himanshu Singh, Senthil Kumaran, Dipanjan Roy, Sourav Banerjee

## Abstract

A high-fat diet (HFD) regulates the feeding circuitry in young-adult mice by stimulating neurogenesis from β2 tanycytes in the hypothalamus. However, the fate of HFD-induced tanycytic neurogenesis, the functional attributes of the nascent neurons, and mechanisms underlying their subsequent integration into the feeding-related neural circuitry remain unclear. microRNAs (miRNAs) are known to play crucial roles in adult neurogenesis. In this study, miRNA profiling of BrdU-labelled neurogenic β2 tanycytes from HFD-fed mice identified a cohort of miRNAs that originate from different chromosomal loci rather than a single genomic cluster. Network analysis of predicted targets for these HFD-induced miRNAs identified key "hub" mRNA targets that influence neurogenesis and the functional integration of newborn neurons. These HFD-induced miRNAs promoted the differentiation of nascent neurons into orexigenic AgRP+ and anorexigenic POMC^+^ neurons. Interestingly, only AgRP+ newborn neurons were selectively integrated into the functional feeding circuit, as detected by c-Fos immunostaining. This integration was disrupted when HFD-induced miRNA activity was suppressed via hypothalamic miRNA sponge expression. Furthermore, HFD feeding increased body weight in female mice, which was abrogated upon the inhibition of HFD-induced miRNAs. Magnetic resonance imaging (MRI) revealed elevated liver fat in HFD-fed mice, which reverted to normal levels when the HFD-induced miRNAs in hypothalamic tanycytes were blocked. Collectively, the findings from our study highlight a miRNA-mediated paradigm that links HFD-induced neurogenesis from β2 tanycytes to weight gain and liver fat accumulation, and identifies the selective integration of AgRP+ neurons into the feeding circuitry as a possible causality of HFD-induced changes to body weight.

**Significance Statement:** This study highlights a miRNA-network that biases the transcriptional program in hypothalamic tanycytes to promote neurogenesis in young adult mice following their exposure to an acute high-fat diet (HFD) paradigm. These HFD-induced miRNAs have the potential to alter the brain’s feeding circuitry by promoting neurogenesis and the preferential functional integration of nascent orexigenic (AgRP+) neurons; resulting in weight gain and hepatic fat accumulation. Collective inhibition of this miRNA-network prevents the integration of newborn AgRP+ neurons and reverses the metabolic consequences of high-fat diet. Our study provides an unexplored link between diet, neurogenesis, and remodeling of the feeding circuitry; and reveals potential therapeutic targets for combating hyperphagic obesity.

## Introduction

Research on adult neurogenesis has extended beyond the classical regions of the subventricular zone and the subgranular zone to include roles of other neurogenic niches in the adult mouse brain (1–3). Particularly, there has been a growing interest in hypothalamic neurogenic niches; their responsiveness to dietary factors further underscores their critical role in metabolic and physiological regulation. Hypothalamic adult neurogenesis in response to acute or sustained dietary changes modulate long-term physiological responses in the body such as aberrant body fat regulation and obesity. Studies show that adult neurogenesis from different niches of the brain occur in concomitance with weight gain in mice exposed to both acute and chronic high fat diet (HFD), though the nature of obesity differs between these feeding paradigms (4, 5). Chronic HFD consumption results in severe obesity (6, 7), whereas acute HFD intake induces reversible weight gain (8).

Acute HFD feeding stimulates neurogenesis in the hypothalamic median eminence (MEm) of adult mice (5). This region contains an active neurogenic niche marked by the presence of Nestin+, radial glia-like ependymal cells called tanycytes (9). Tanycytes are metabolite sensing ependymal cells lining the ventral third ventricular wall; anatomically characterized into α1-, α2-, β1-, and β2-subtypes. They are capable of sensing metabolites from the cerebrospinal fluid and blood due to their position outside the blood-brain barrier and have characteristically long projections to the dorsomedial hypothalamus, ventromedial hypothalamus, arcuate nucleus (ArcN) and the median eminence (MEm) (9). Neural circuits underlying feeding behaviour in the mouse brain is centralized to the ArcN, which integrates opposing feeding signals due to the presence of two neuron types: anorexigenic POMC+ neurons promoting satiety, and orexigenic AgRP+ neurons that drive hunger (10–12). AgRP+ and POMC+ neurons are dimorphic and they reciprocally inhibit each other (12). β1-and β2-tanycytes relay signals to activate the ArcN feeding circuits (13, 14) and therefore influence the function of both AgRP+ and POMC+ neurons. β2-tanycytes are Sox2+/Nestin+/Vimentin+ diet-responsive neural progenitors that adapt to a neural fate in response to variations in dietary factors (5). Neurogenesis from β2-tanycytes is critical for HFD-induced metabolic adaptation in female mice; previous studies report that irradiation of the MEm ablates β2-tanycytes and prevents HFD-driven weight gain (5). However, how HFD-induced neurogenesis occurs from β2-tanycytes and how these nascent neurons are integrated into the feeding circuit is unknown.

Among the various effectors influencing diet-induced modulation of the hypothalamic feeding circuits, miRNAs play a unique but understudied role. These small noncoding RNAs modulate mRNA translation and RNA stability in response to external cues, positioning them as likely regulators of metabolic pathways during both acute and chronic dietary changes. Studies confirm that hypothalamic miRNA expression changes due to both acute and chronic nutritional interventions in rats (15, 16), with specific miRNAs (let-7a, miR-9, miR-30e, miR-103, miR-132, miR-145, miR-200a, miR-218) responding to either calorie restriction or HFD. Notably, administration of miR-103-mimic to the ArcN of Dicer KO mice attenuated hyperphagic obesity (16). miRNAs are also established regulators of adult neurogenesis from the subventricular zone of the lateral ventricle and the subgranular zone of the hippocampus (17, 18). Thus, miRNAs likely mediate HFD-induced tanycytic neurogenesis and subsequent circuit modifications (18–23), bridging dietary inputs to neural adaptations.

In this study, we profiled miRNAs in BrdU+ β2-tanycytes from female C57BL/6j mice after acute HFD exposure, identifying a cohort of diet-induced miRNAs. By conducting a protein–protein interaction network analysis for proteins encoded by mRNAs that are targeted by this collective cohort of miRNAs, we uncovered a miRNA-mediated molecular framework involved in HFD-induced neurogenesis from β2-tanycytes. Fate-mapping demonstrated differentiation of the nascent neurons into both orexigenic AgRP+ and anorexigenic POMC+ types, but only AgRP+ neurons showed functional circuit integration as evidenced by c-Fos expression. Inhibition of HFD-induced miRNAs in the hypothalamus disrupted this neuronal integration, which in turn reversed HFD-induced body weight gain. Magnetic resonance imaging (MRI) indicates that suppressing these miRNAs also inhibited the accumulation of liver fat. Our findings unveil an unexplored miRNA-dependent mechanism regulating body weight through hypothalamic neurogenesis and selective orexigenic neuron recruitment into the feeding circuit in female mice following acute exposure to a high-fat diet.

## Results

### HFD induces the upregulation of a cohort of miRNAs in neurogenic β2 tanycytes

To identify diet-regulated miRNAs from neurogenic tanycytes present in the median eminence (MEm), female mice were either given normal chow diet (NCD) or HFD during P45 – P75 and tanycytes undergoing neurogenesis were labelled by BrdU incorporation (Fig. 1A). Female mice were specifically used for this study as previous research has demonstrated that the median eminence of female, but not male, mice showed enhanced neurogenesis after HFD feeding (8). Using laser capture microscopy, BrdU^+^ tanycytes (Fig. 1B) were isolated from the median eminence and the differential expression of miRNAs were analyzed by miRNA arrays. We observed that 32 miRNAs were differentially expressed upon HFD feeding, whereas 5 miRNAs were differentially expressed in NCD fed mice. Among these transcripts, a cohort of miRNAs; miR-382, miR-145, miR-196, miR-1894 and let-7b (Table 1) were significantly up-regulated upon HFD feeding (Fig. 1C); whereas NCD feeding significantly enhanced the expressions of let-7d, let-7i and let-7e (Table 1) (Fig. 1C). Interestingly, these miRNAs are not localized to a single genomic cluster, rather they originate from different chromosomes (Fig. 1D). Our results indicate that miRNAs are differentially expressed in the neurogenic β2-tanycytes of female mice following acute exposure to HFD.

**Fig. 1:**
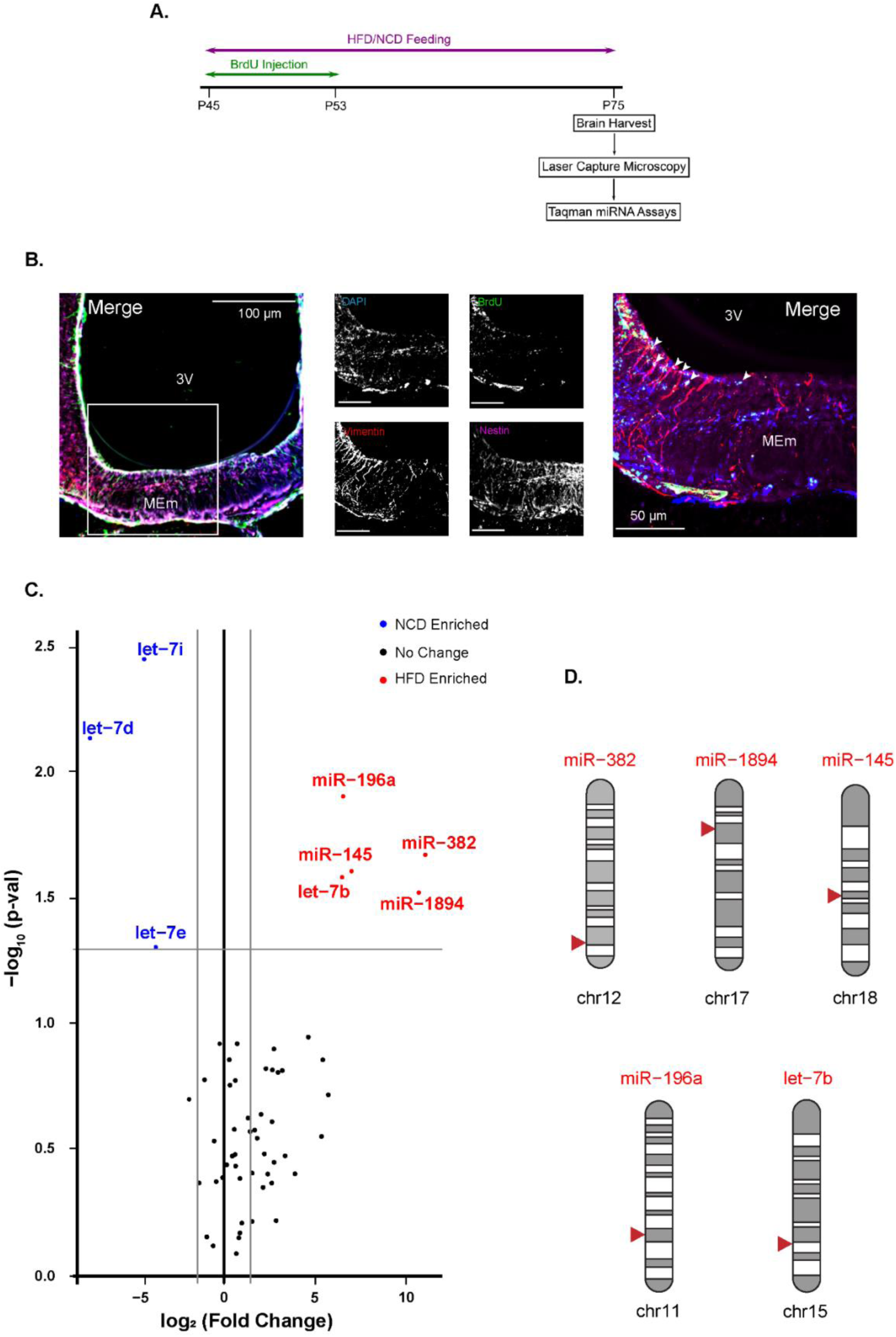
Differentially expressed miRNAs identified from adult neurogenic tanycytes in median eminence. (A) Experimental paradigm to determine the miRNA expression profiles by isolation of BrdU labelled β2 neurogenic tanycytes of the median eminence. (B) Photomicrograph showing BrdU labelling of Nestin^+^ and Vimentin^+^ β2 neurogenic tanycytes in median eminence (MEm) post 30 days of HFD feeding. (C) Scatter volcano plot showing expression of miRNAs from β2 neurogenic tanycytes following HFD (red) or NCD (blue) feeding. Significant changes in miRNA expression (marked by arrow). N = 3 independent experiments. Two-tailed t-test was performed and post hoc false discovery rates (FDR) was analyzed using the Benjamini-Hochberg method. ArcN: Arcuate Nucleus, 3V: Third ventricle, MEm: Median Eminence. Scale bars are as indicated. (D) Karyotype Phenogram shows the genomic locations of differentially expressed miRNAs.

**Table 1:**
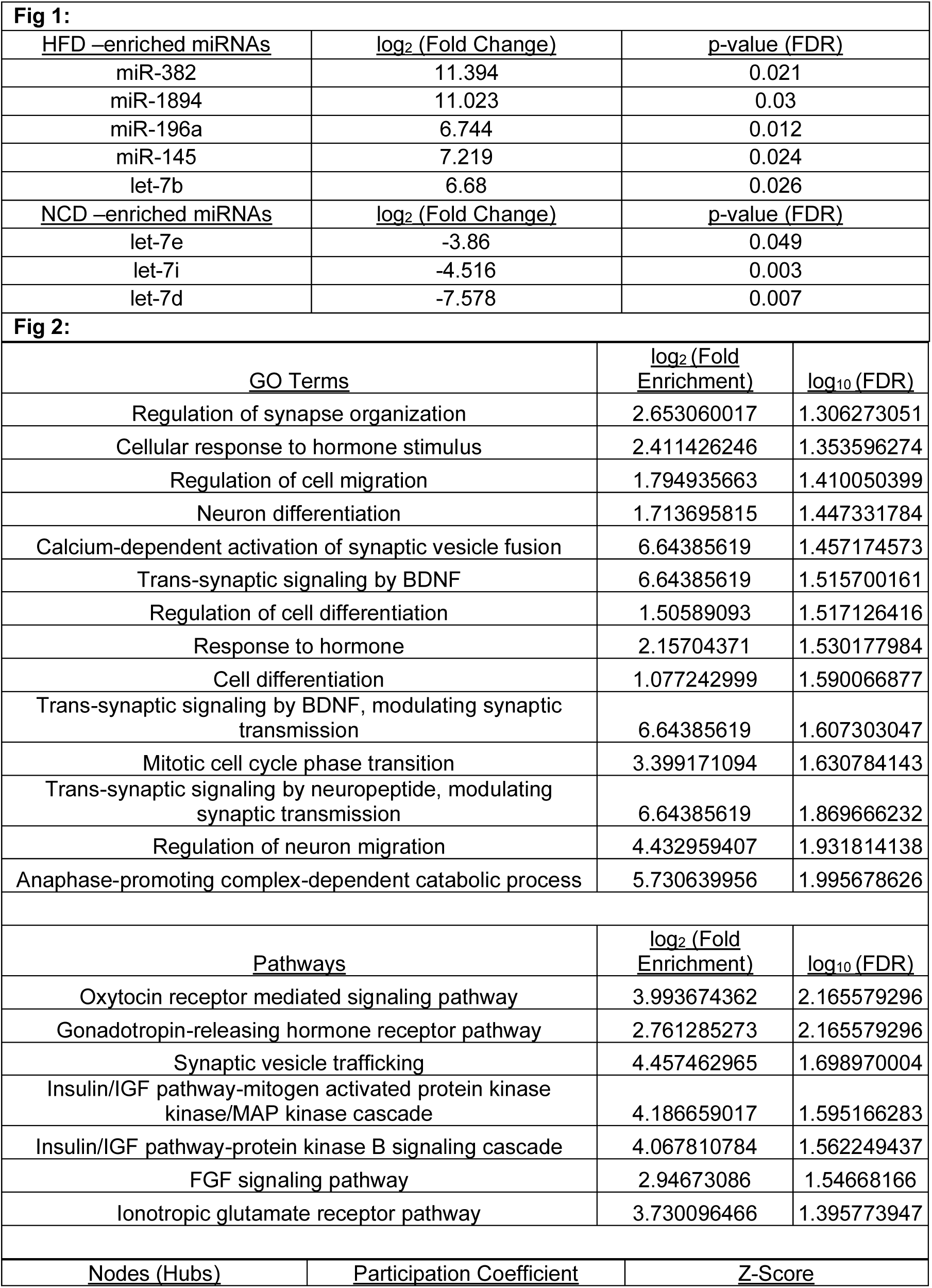
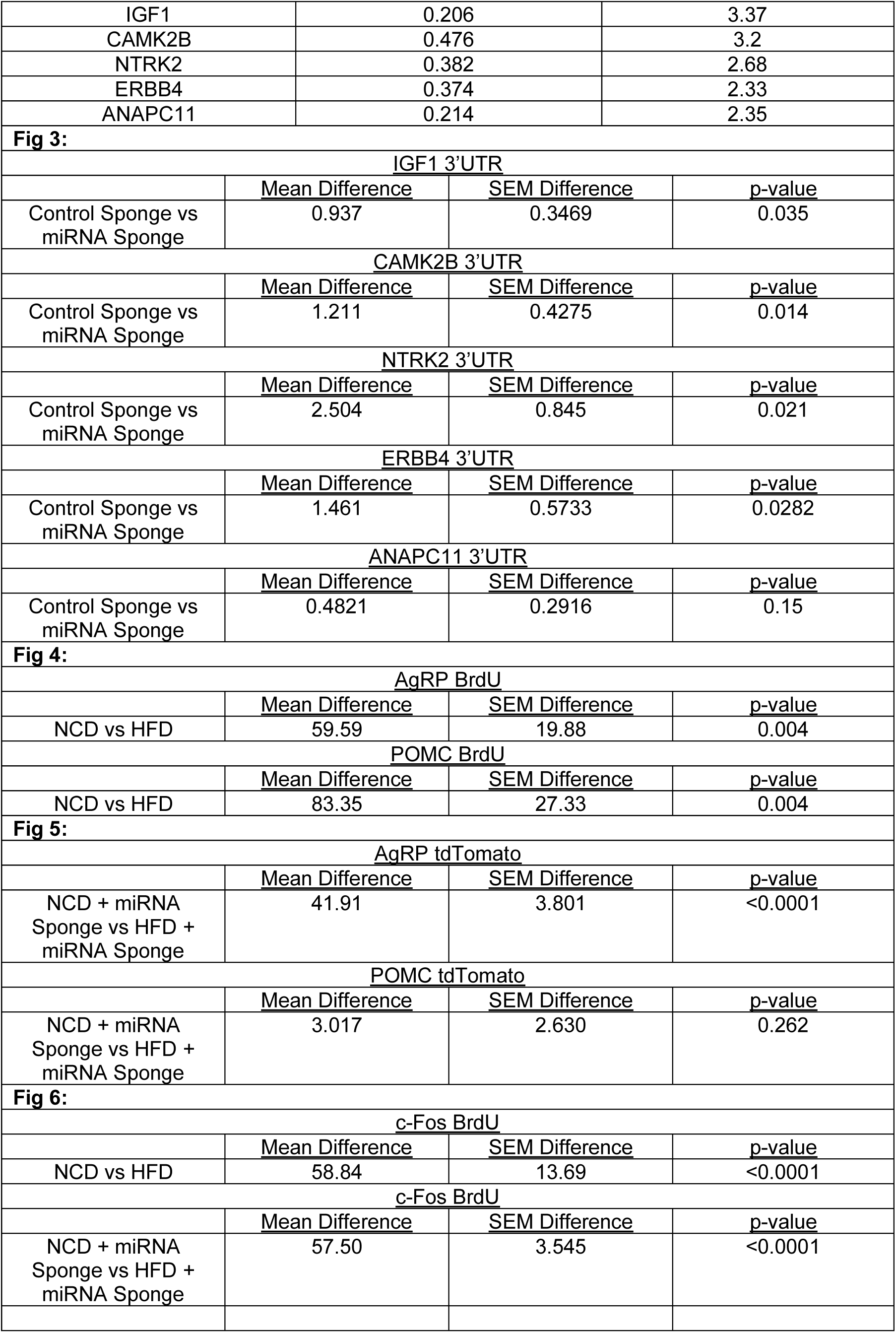
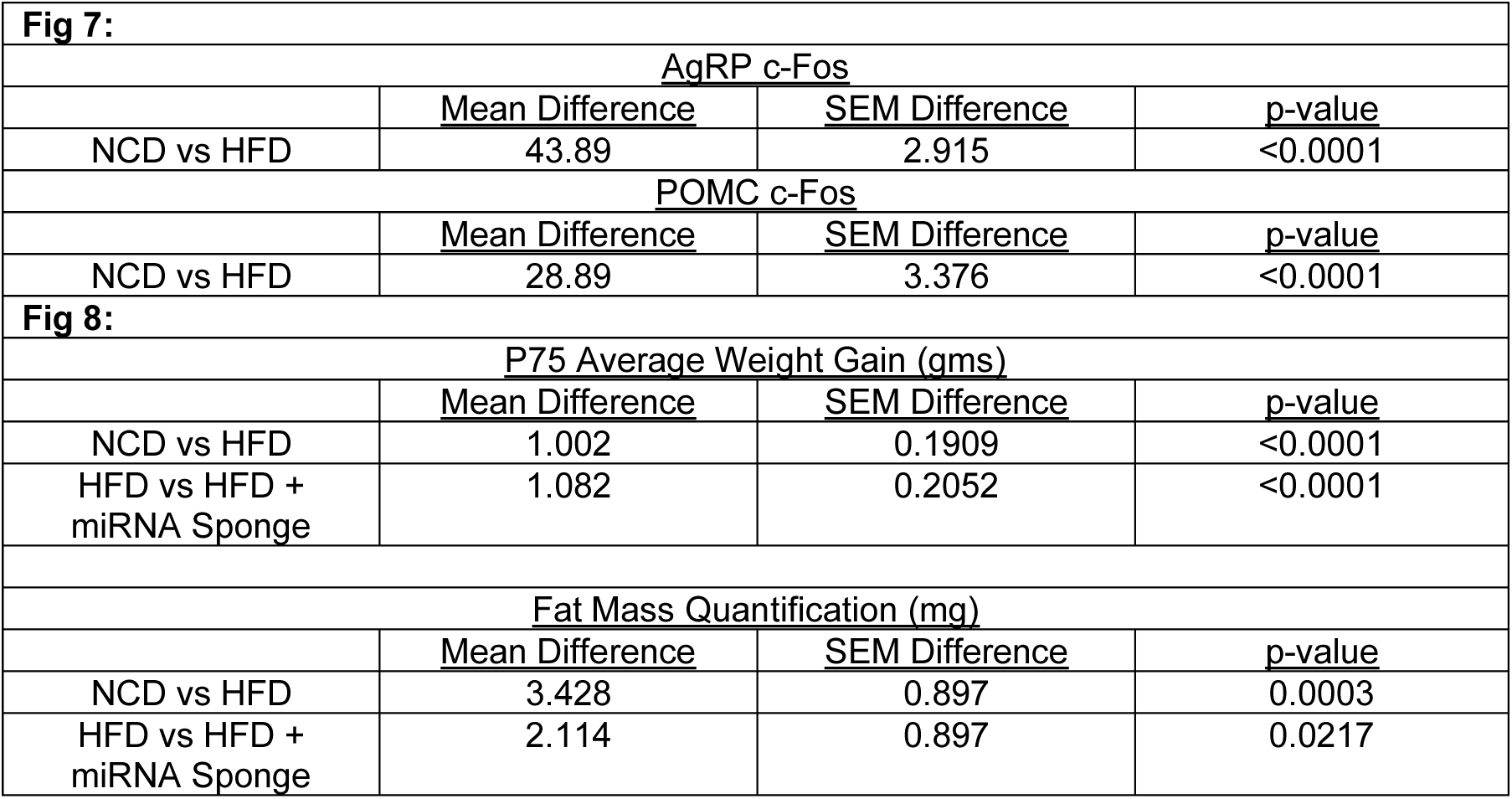
Data.

### Diet-induced miRNAs regulate a network of transcripts involved in neurogenesis

To determine how miRNAs regulate the neurogenesis from β2 tanycytes, mRNA targets of the HFD-responsive miRNAs; miR-1894, miR-382, miR-145, miR-196 and let-7b were analyzed. We reasoned that since miRNAs exert regulatory control on the translation or stability of mRNAs, the mRNAs that would be most affected due to HFD are likely to possess binding sites for all five HFD-induced miRNAs. Accordingly, putative target mRNAs containing binding sites for all five miRNAs were obtained from the TargetScan, miRdb and PicTar databases. We have selected 1204 transcripts represented in all three databases and analyzed the interactions between them using StringDB. Graph theory-based network analysis was performed to determine the key miRNA target transcripts. Nodes (mRNA targets) were clustered using Louvain’s community clustering method (24). Stable optimization of the clustering of the nodes using the above method was achieved using a resolution parameter ranging from 0.5 – 0.75 (Fig. 2A). We observed a highly modular network among the targets of the upregulated miRNAs upon HFD feeding, whereas this modularity is absent upon NCD feeding. We have detected 11 modules in the HFD network, out of which 6 modules are highlighted (Fig. 2B). This network modularity is a key target selection criterion for further analysis as clustered transcripts within a network have the potential to work concertedly to drive neurogenesis in the tanycytes and subsequent neuronal differentiation upon HFD feeding. Among the modularized community clusters, inter-community and intra-community measures of nodal activity were analyzed to determine the important target nodes that are critical for the entire network. Participation coefficient, a marker for inter-community interactions, was analyzed for all nodes to define the hubs. We have analyzed z-scores, a marker for intra-community interactions, to select the critical hubs in the network. Hubs with z-scores > 2 were selected for Gene Ontology (GO) analysis. We reasoned that these hubs contain key transcripts that can regulate multiple nodes. mRNA targets that are positioned at the “hubs” of the network, are considered to be central effectors of diet induced neurogenesis and neuronal differentiation. GO analysis of the mRNA targets revealed significantly enriched GO-terms including neuron differentiation, mitotic cell cycle phase transition, cellular response to hormone stimulus, trans-synaptic signaling by BDNF, trans-synaptic signaling by neuropeptide modulating synaptic transmission, regulation of cell migration and anaphase-promoting complex-dependent catabolic (Table 1) (Fig. 2C). We have also assessed significant GO-enriched pathways that includes insulin/IGF pathway-protein kinase B signaling cascade, insulin/IGF pathway-mitogen activated protein kinase cascade, ionotropic glutamate receptor pathway, FGF-signaling pathway and gonadotropin-releasing hormone receptor pathway (Table 1) (Fig. 2D). Based on the GO-enrichment analysis, we have selected IGF1, NTRK2, CAMK2B, ERBB4 and ANAPC11 (Table 1) for further validation *in vitro* using a reporter assay (Fig. 2E).

**Fig. 2:**
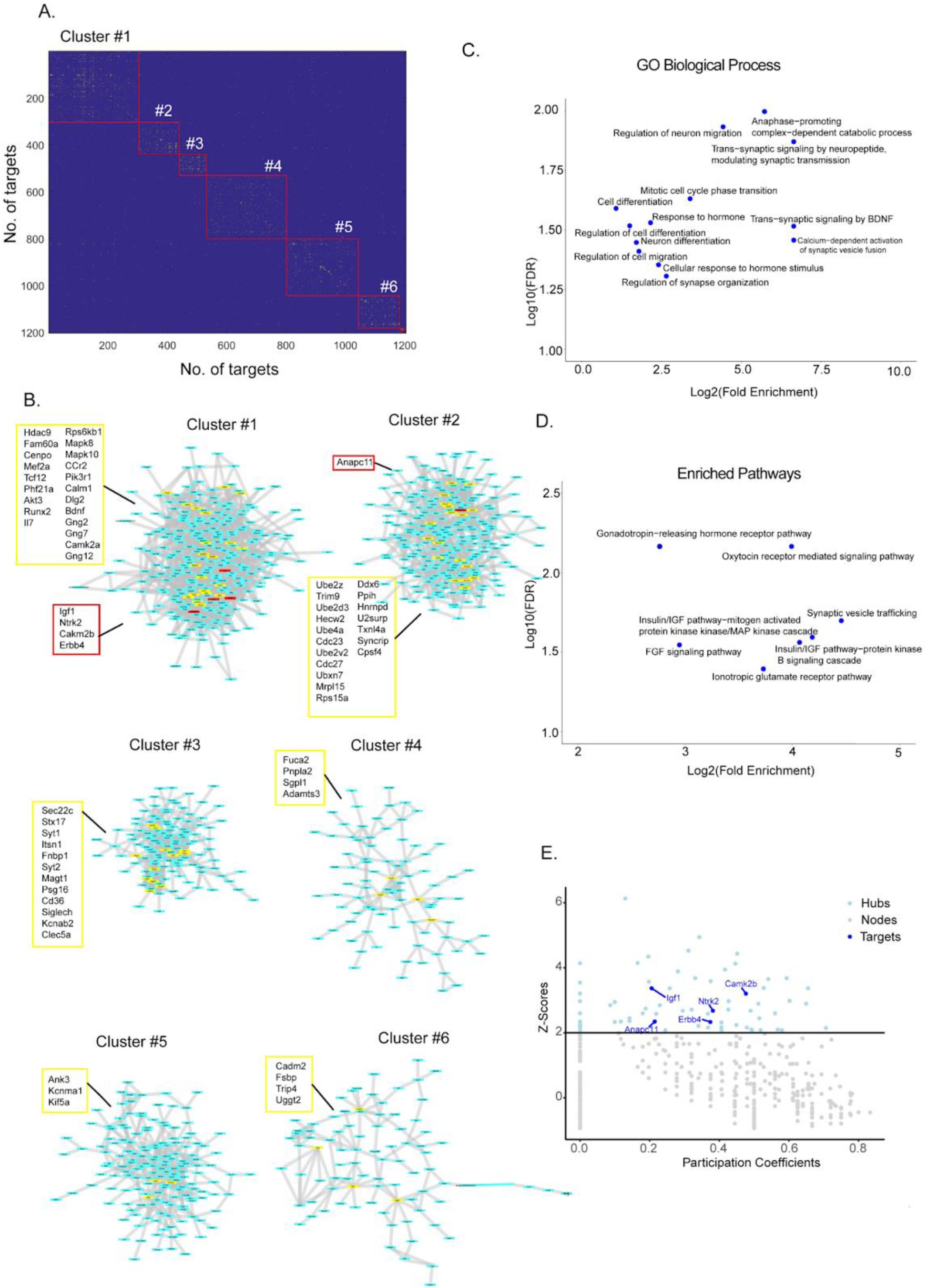
Weighted gene coexpression network analysis of target transcripts containing binding sites of HFD-regulated miRNAs. (A) Adjacency matrix showing community clustering (red boxes) based on the interaction between each target as a node. (B). 6 clusters of the network of target transcripts containing binding sites for all five miRNAs. All hubs are indicated in the “yellow” box. Hubs selected for target validation via reporter assay are represented in the “red” box. (C) Scatter plots showing fold enrichment of significant GO-terms in Biological Process. (D) Scatter plots showing fold enrichment of significant GO-terms from pathway analysis. (E) Scatter plot showing the z-scores and the participation coefficients for all nodes in the network. Nodes are considered “hubs” for z-score > 2. Hubs selected for reporter analysis are indicated.

### HFD-induced miRNAs directly bind to the 3’ UTR of mRNAs that are the central effectors of neurogenesis

To experimentally validate the interactions between HFD-regulated miRNAs and their “hub” mRNA targets identified in our network analysis, we performed a luciferase reporter assay following the inhibition of all five HFD-induced miRNAs. The key hub transcripts IGF1, NTRK2, CAMK2B, ERBB4 and ANAPC11 were selected for the validation of miRNA-mRNA interactions *in vitro*; owing to their previous implications in neurogenesis (IGF1), neuronal development (ANAPC11, IGF1), synaptogenesis (CAMK2B,ERBB4), energy homeostasis (NTRK2) or hyperphagic obesity (NTRK2) (25–36).

To sequester five miRNAs simultaneously, a miRNA sponge (containing complementary binding sites to all five miRNAs) was cloned into an AAV vector that co-expressed tdTomato (Fig. 3A). The efficacy of the miRNA sponge was determined by the fluorescence intensity of tdTomato expression. Administration of the miRNA sponge significantly reduced the intensity of tdTomato expression (65.05% ± 12.52% decrease, p<0.0001) (Fig. S1B-S1C) as compared to that in the control vector. The 3’UTRs of each target mRNA containing the binding sites of all five miRNAs were fused to the Gaussia luciferase reporter (Fig. 3B). We co-transfected the miRNA sponge along with each of the reporters in Neuro-2a cells. Simultaneous inhibition of all five HFD-induced miRNAs led to the significant enhancement of luciferase reporter activity driven by the 3’UTRs of IGF1, NTRK2, CAMK2B and ERBB4 (Fig. 3C); but not ANAPC11 (Table 1) (Fig. 3C).

**Fig. 3:**
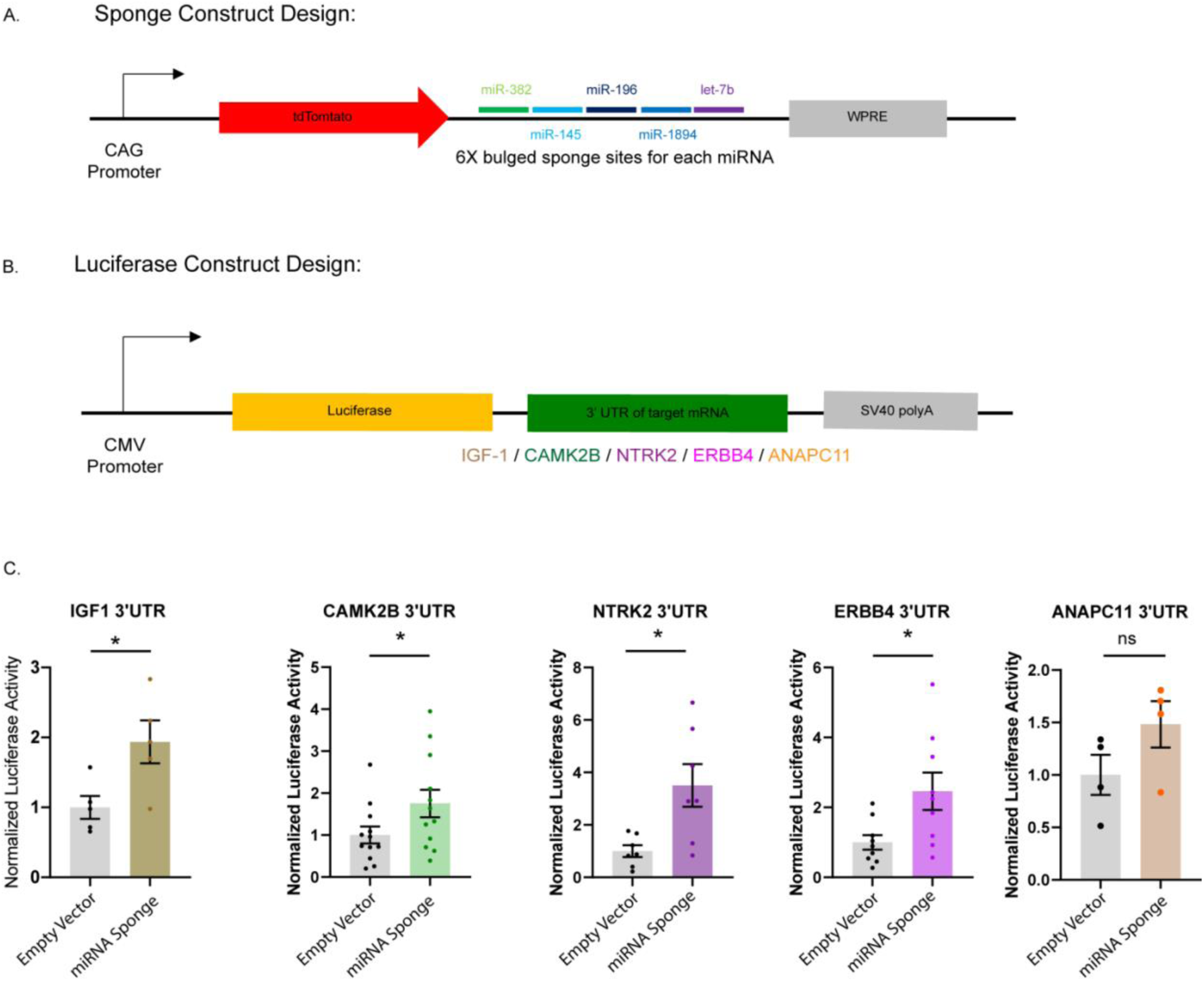
*In vitro* validation of mRNA target interactions with HFD responsive miRNAs. (A) Schematic design of miRNA sponge construct. (B) Schematic design of luciferase reporter. (C) Quantitation of luciferase reporter expression driven by 3’UTRs of IGF1, NTRK2, CAMK2B, ERBB4 and ANAPC11 following simultaneous inhibition of all 5 HFD-regulated miRNAs’ activity. N = 4 – 11 independent experiments. *p < 0.05. Data shown as mean ± SEM. Two tailed t-test with Welch’s correction.

### HFD-induced miRNAs specifically promote the generation of nascent AgRP^+^ neurons

To determine the fate of adult born BrdU^+^ nascent neurons upon HFD feeding, we have analyzed whether these neurons differentiate into POMC-expressing anorexigenic or AgRP-expressing orexigenic neurons in the MEm of the hypothalamus. Following HFD/NCD feeding, BrdU labelled cells were analyzed for the co-expression of AgRP and POMC along with NeuN. HFD-fed mice showed a significant enhancement of AgRP expressing BrdU^+^ neurons (59.59% ± 19.88% increase, p = 0.004) as compared to mice fed with NCD (Fig. 4A and Fig. 4B). We detected a similar increase of POMC expressing BrdU^+^ neurons (83.35% ± 27.33% increase, p = 0.004) in HFD-fed mice (Fig. 4C – 4D). Next, we analyzed whether inhibition of the HFD-induced miRNAs could affect the generation of AgRP^+^ or POMC^+^ neurons. A miRNA sponge against all the five HFD-upregulated miRNAs was cloned into AAV-CAG-tdTomato and the AAV-tdTomato-miRNA sponge virus injected into the hypothalamus of adult female mice. miRNA sponge expression was confirmed by the expression of the tdTomato reporter (Fig. S1). The resultant inhibition of the HFD-induced miRNAs reduced the number of AgRP^+^ neurons (43.89% ± 2.915% decrease, p < 0.0001) (Fig. 5A – 5B). In contrast, there was a minimal but statistically significant increase in POMC^+^ neurons (28.89% ± 3.376% increase, p < 0.0001) (Fig. 5C – 5D). Our observations reveal a surprising insight: acute HFD-feeding paradigm selectively promotes the generation of AgRP+ orexigenic neurons through a miRNA-dependent mechanism, while the generation of POMC+ anorexigenic neurons remains unaffected by these miRNAs.

**Fig. 4:**
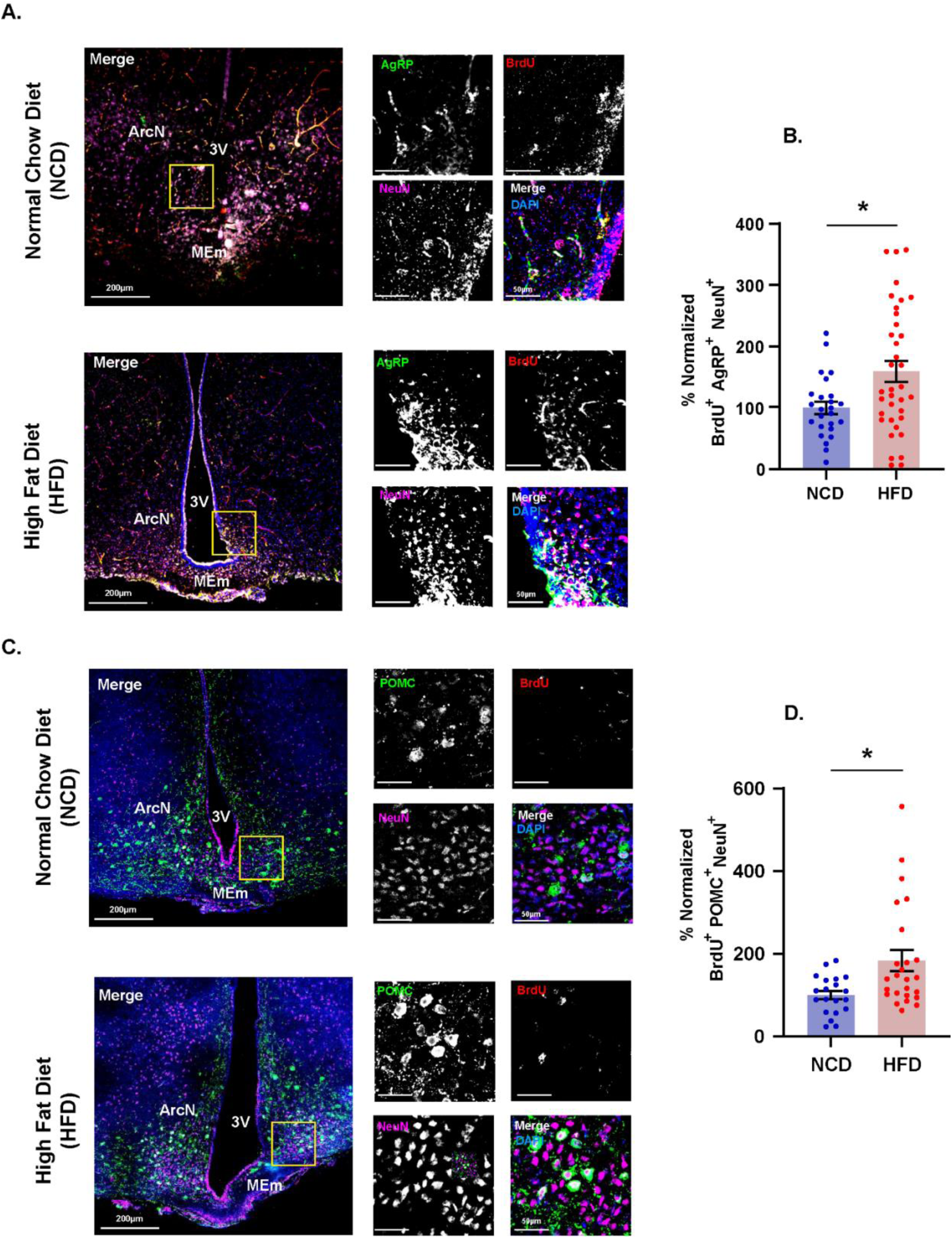
Cell fate of tanycytic derived neurons in the hypothalamic feeding circuitry. (A) Photomicrographs represent orexigenic fate of diet-induced nascent neurons upon HFD or NCD feeding. Low magnification image of brain section of all channels [DAPI (blue), BrdU (red), AgRP (green), NeuN (magenta)] were merged (white). High magnification images of brain sections labelled with DAPI (blue), BrdU (red), AgRP (green), NeuN (magenta) and merged (white). Scale bars are as indicated. (B) Quantitative analysis of the number of colocalized BrdU^+^ cells with AgRP^+^ and NeuN^+^ in the feeding circuit of mice after NCD and HFD feeding. n = 25-36 sections. N = 3 mice. *p < 0.005. Data shown as mean ± SEM. Two tailed t-test with Welch’s correction. (C) Photomicrographs represent anorexigenic fate of diet-induced nascent neurons upon HFD or NCD feeding. Low magnification image of brain section of all channels [DAPI (blue), BrdU (red), POMC (green), NeuN (magenta)] were merged (white). High magnification images of brain sections labelled with DAPI (blue), BrdU (red), POMC (green), NeuN (magenta) and merged (white). Scale bars are as indicated. (B) Quantitative analysis of the number of colocalized BrdU^+^ cells with POMC^+^ and NeuN^+^ in the feeding circuit of mice after NCD and HFD feeding. n = 20-25 sections. N = 3 mice. *p < 0.005. Data shown as mean ± SEM. Two tailed t-test with Welch’s correction.

**Fig. 5:**
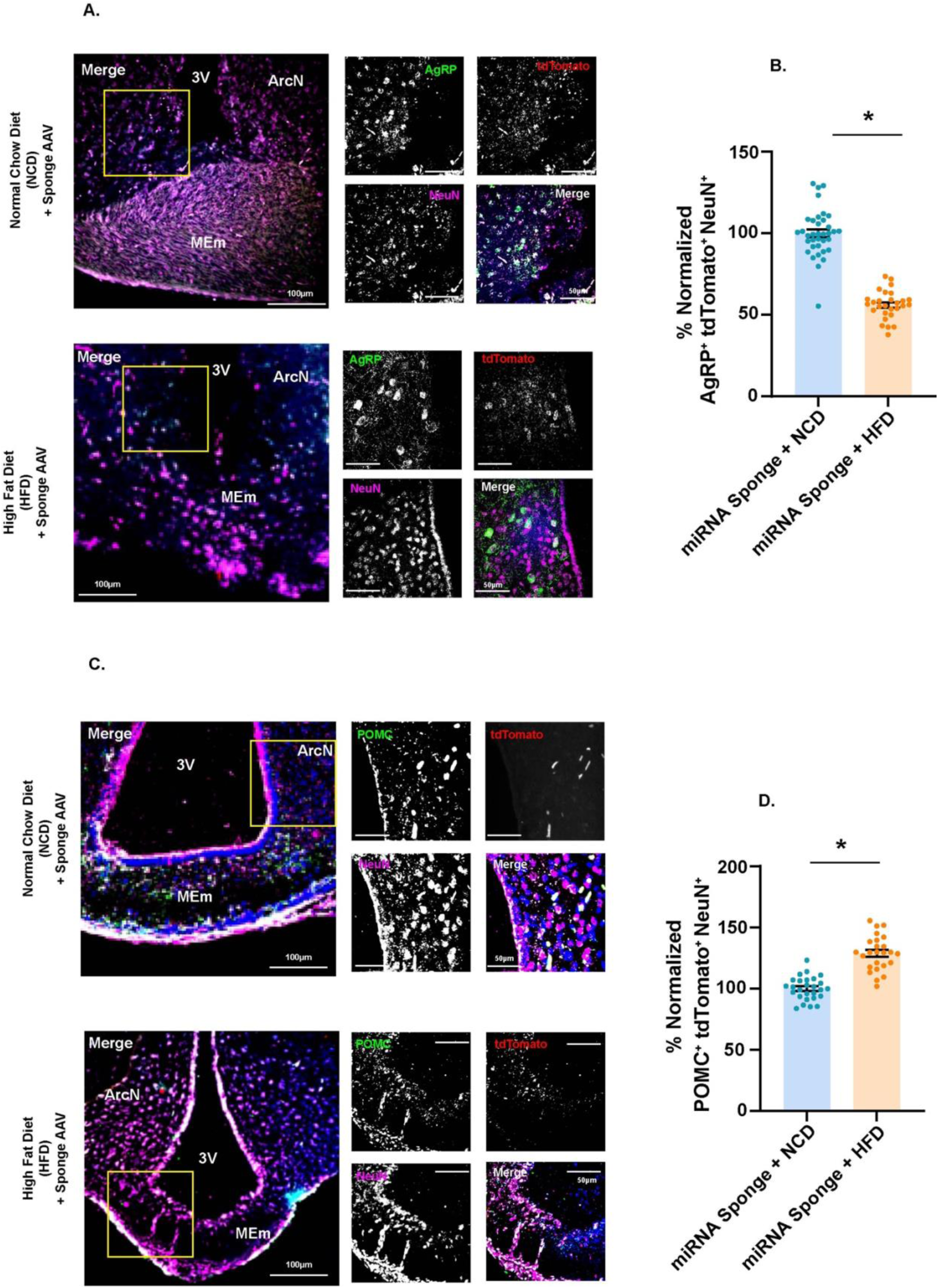
Inhibition of HFD –induced miRNAs prevents generation of orexigenic AgRP neurons. (A) Photomicrographs represent abrogation of orexigenic fate of the hypothalamic feeding circuit upon HFD or NCD feeding. Low magnification image of brain section of all channels [DAPI (blue), AAV –expressed tdTomato (red), AgRP (green), NeuN (magenta)] were merged (white). High magnification images of brain sections labelled with DAPI (blue), tdTomato (red), AgRP (green), NeuN (magenta) and merged (white). Scale bars are as indicated. (B) Quantitative analysis of the number of colocalized AgRP^+^ cells with tdTomato^+^ and NeuN^+^ in the feeding circuit of mice after NCD and HFD feeding. n = 28-36 sections. N = 3 mice. *p < 0.0001. Data shown as mean ± SEM. Two tailed t-test with Welch’s correction. (C) Photomicrographs represent increase of anorexigenic fate of the hypothalamic feeding circuit upon HFD or NCD feeding. Low magnification image of brain section of all channels [DAPI (blue), AAV –expressed tdTomato (red), POMC (green), NeuN (magenta)] were merged (white). High magnification images of brain sections labelled with DAPI (blue), tdTomato (red), POMC (green), NeuN (magenta) and merged (white). Scale bars are as indicated. (D) Quantitative analysis of the number of colocalized POMC^+^ cells with tdTomato^+^ and NeuN^+^ in the feeding circuit of mice after NCD and HFD feeding. n = 25-27 sections. N = 3 mice. *p < 0.0001. Data shown as mean ± SEM. Two tailed t-test with Welch’s correction.

### HFD-induced miRNAs promote the functional integration of newborn orexigenic AgRP**^+^** neurons

We have quantified the number of BrdU^+^ and c-Fos^+^ neurons in the MEm and ArcN adjacent to MEm as these areas contain mature POMC and AgRP neurons. AAVs containing either AAV-tdTomato-miRNA sponge vector or control vector was injected into adult female mice and the number of BrdU+ and c-Fos+ neurons were measured with or without the inhibition of HFD-induced miRNAs. We observed a significant increase in BrdU^+^ and c-Fos^+^ neurons upon HFD feeding (Table 1) (Fig. 6A - 6B) as compared to mice fed with NCD. The simultaneous inhibition of all HFD-induced miRNAs resulted in a significant decrease in BrdU^+^ and c-Fos^+^ neurons (Table 1) (Fig. 6C and 6D). Our data demonstrates that inhibition of HFD-induced miRNAs precludes the formation of nascent neurons with functional properties.

**Fig. 6:**
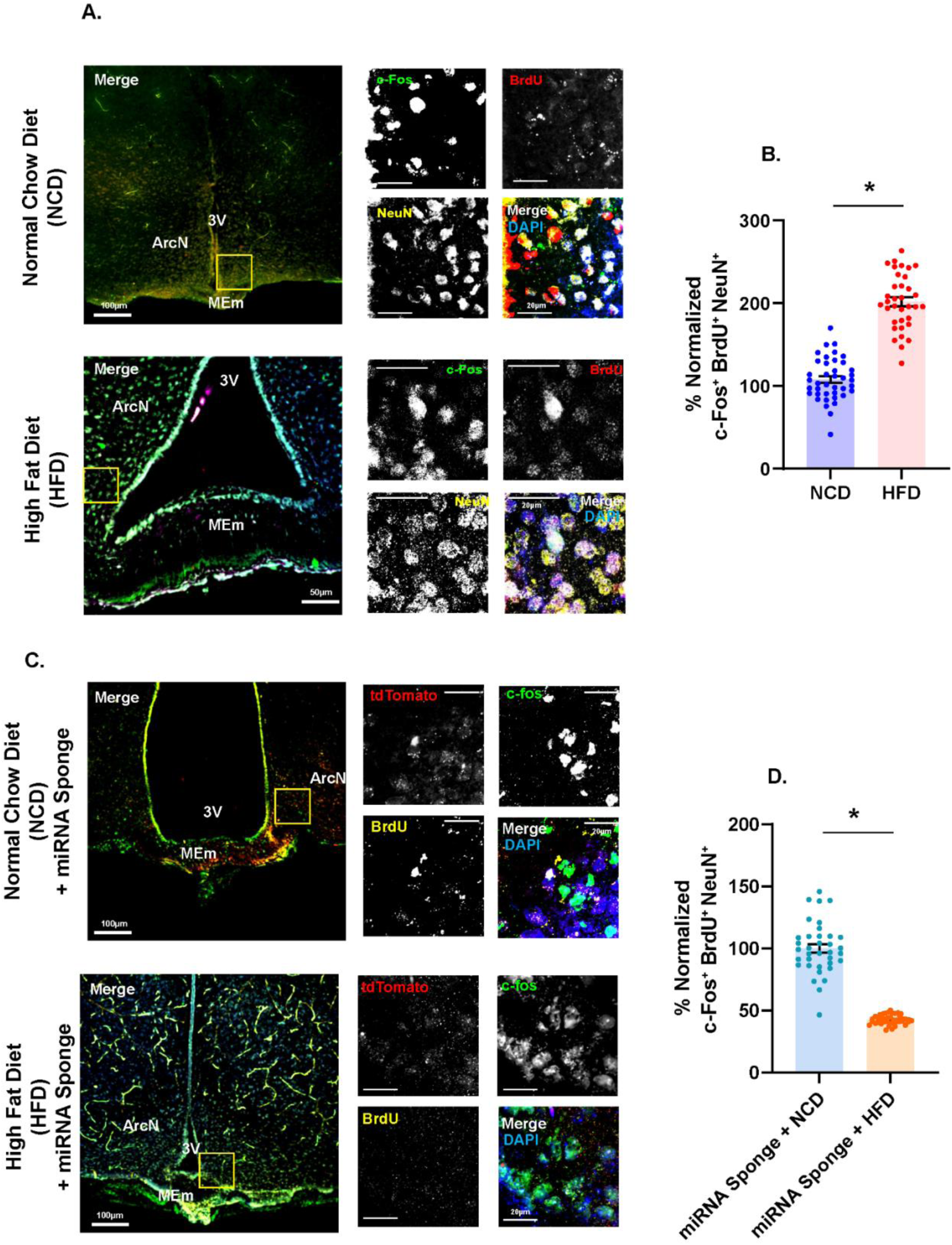
Inhibition of HFD -induced miRNAs block the activity of newborn neurons. (A) Photomicrographs represent functional activity of diet-induced nascent neurons in the hypothalamic feeding circuitry upon HFD or NCD feeding. Low magnification image of brain sections of all channels [DAPI (blue), BrdU (red), c-Fos (green), NeuN (yellow)] were merged (white). High magnification images of brain sections labelled with DAPI (blue), BrdU (red), c-Fos (green), NeuN (yellow) and merged (white). Scale bars are as indicated. (B) Quantitative analysis of the number of colocalized BrdU^+^ cells with c-Fos^+^ and NeuN^+^ in the feeding circuit of hypothalamus after NCD and HFD feeding. n = 35-40 sections. N = 3 mice. *p < 0.0001. Data shown as mean ± SEM. Two tailed t-test with Welch’s correction. (C) Photomicrographs represent abrogation of functional activity of diet-induced nascent neurons upon HFD or NCD feeding in the hypothalamic feeding circuitry. Low magnification image of brain sections of all channels [DAPI (blue), AAV –expressed tdTomato (red), c-Fos (green), NeuN (yellow)] were merged (white). High magnification images of brain sections labelled with DAPI (blue), tdTomato (red), c-Fos (green), BrdU (yellow) and merged (white). Scale bars are as indicated. (B) Quantitative analysis of the number of colocalized BrdU^+^ cells with c-Fos^+^ and tdTomato^+^ in the feeding circuit of mice after NCD and HFD feeding. n = 36-41 sections. N = 3 mice. *p <0.0001. Data shown as mean ± SEM. Two tailed t-test with Welch’s correction.

We then examined the activity of orexigenic AgRP^+^ and anorexigenic POMC^+^ neurons by c-Fos immunostaining after HFD feeding. HFD feeding promoted the activity of AgRP^+^ (41.91% ± 3.801 % increase, p<0.0001), but not POMC^+^ neurons (Fig. 7A – 7D).

**Fig. 7:**
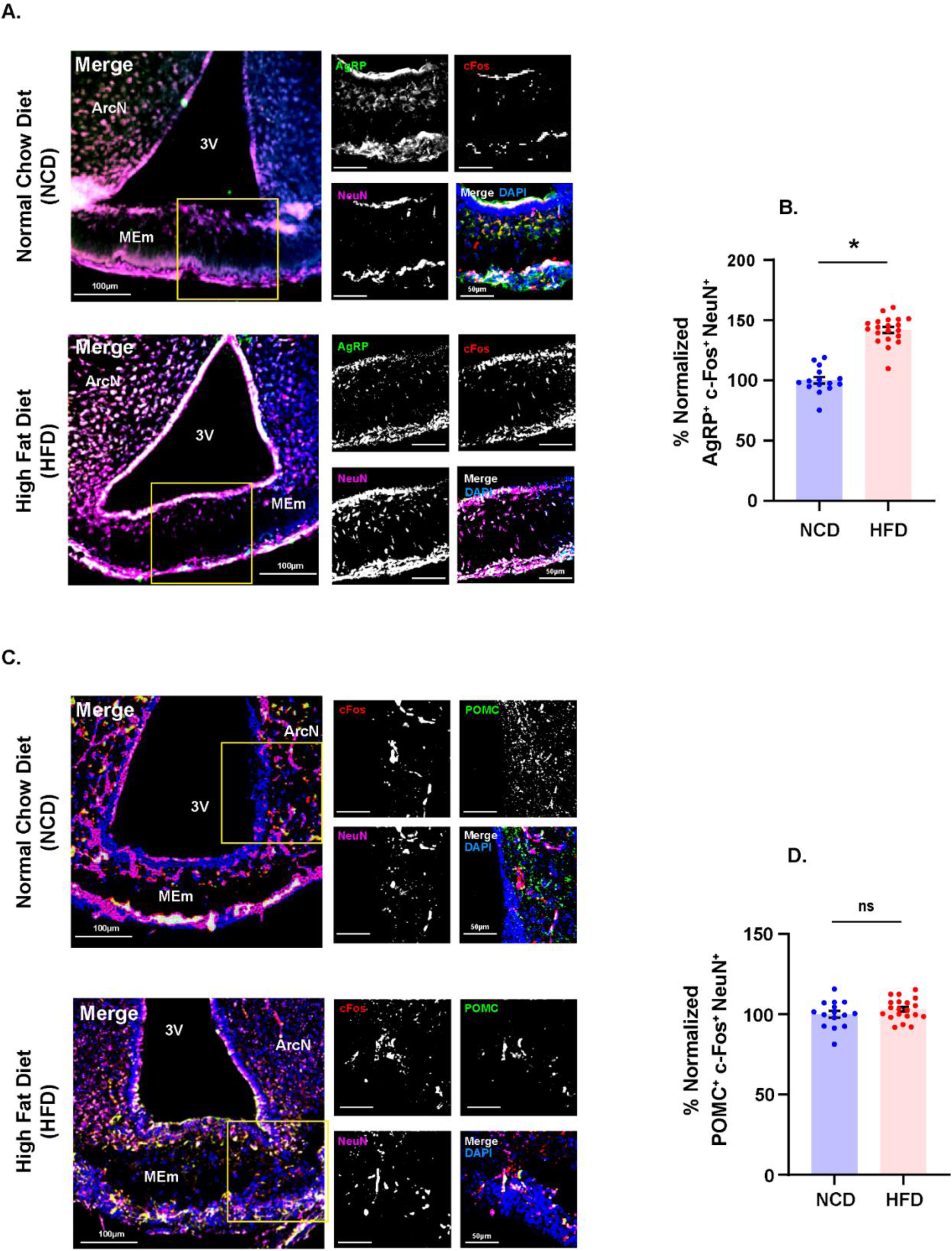
HFD administration enhances the activity of AgRP neurons in hypothalamus. (A) Photomicrographs represent abrogation of functional activity of orexigenic neurons in the hypothalamic feeding circuitry upon HFD or NCD feeding. Low magnification image of brain sections of all channels [DAPI (blue), c-Fos (red), AgRP (green), NeuN (magenta)] were merged (white). High magnification images of brain sections labelled with DAPI (blue), c-Fos (red), AgRP (green), NeuN (magenta) and merged (white). Scale bars are as indicated. (B) Quantitative analysis of the number of colocalized AgRP^+^ cells with c-Fos^+^ and NeuN^+^ in the feeding circuit of hypothalamus after NCD and HFD feeding. n = 15-20 sections. N = 3 mice. *p < 0.0001. Data shown as mean ± SEM. Two tailed t-test with Welch’s correction. (C) Photomicrographs represent functional activity of orexigenic neurons in the hypothalamic feeding circuitry upon HFD or NCD feeding. Low magnification image of brain sections of all channels [DAPI (blue), c-Fos (red), POMC (green), NeuN (magenta)] were merged (white). High magnification images of brain sections labelled with DAPI (blue), c-Fos (red), POMC (green), NeuN (magenta) and merged (white). Scale bars are as indicated. (D) Quantitative analysis of the number of colocalized POMC^+^ cells with c-Fos^+^ and NeuN^+^ in the feeding circuit of hypothalamus after NCD and HFD feeding. n = 15-20 sections. N = 3 mice. p = 0.262. Data shown as mean ± SEM. Two tailed t-test with Welch’s correction.

### Inhibition of HFD-induced miRNAs in hypothalamus prevents body weight gain and the concomitant increase in liver fat mass

We wanted to investigate the physiological relevance of the HFD-feeding induced, miRNA-dependent neurogenesis of orexigenic AgRP^+^ neurons and their integration into the feeding circuitry. Accordingly, we examined alterations in body weight and hepatic lipid accumulation following the inhibition of diet -induced miRNAs in the presence of HFD. AAVs containing the AAV-tdTomato-miRNA sponge vector or control vector was injected into the MEm and the ArcN (adjacent to MEm) of female mice at P25, following which the mice were subjected to HFD / NCD feeding paradigms during P45 - P75 and their body weight measured for the duration. HFD-feeding resulted in enhanced body weight (Table 1) at P75. This gain was abrogated after inhibition of diet-induced miRNAs (Fig. 8A – 8B) despite HFD-feeding. We further used MRI to determine the accumulation of hepatic fat following the inhibition of HFD-induced miRNAs in the presence or absence of HFD. HFD feeding led to enhanced fat mass in the liver (Table 1). The accumulation of liver fat mass in female mice was abolished when HFD-induced miRNAs were inhibited (Fig. 8C – 8D).

**Fig. 8:**
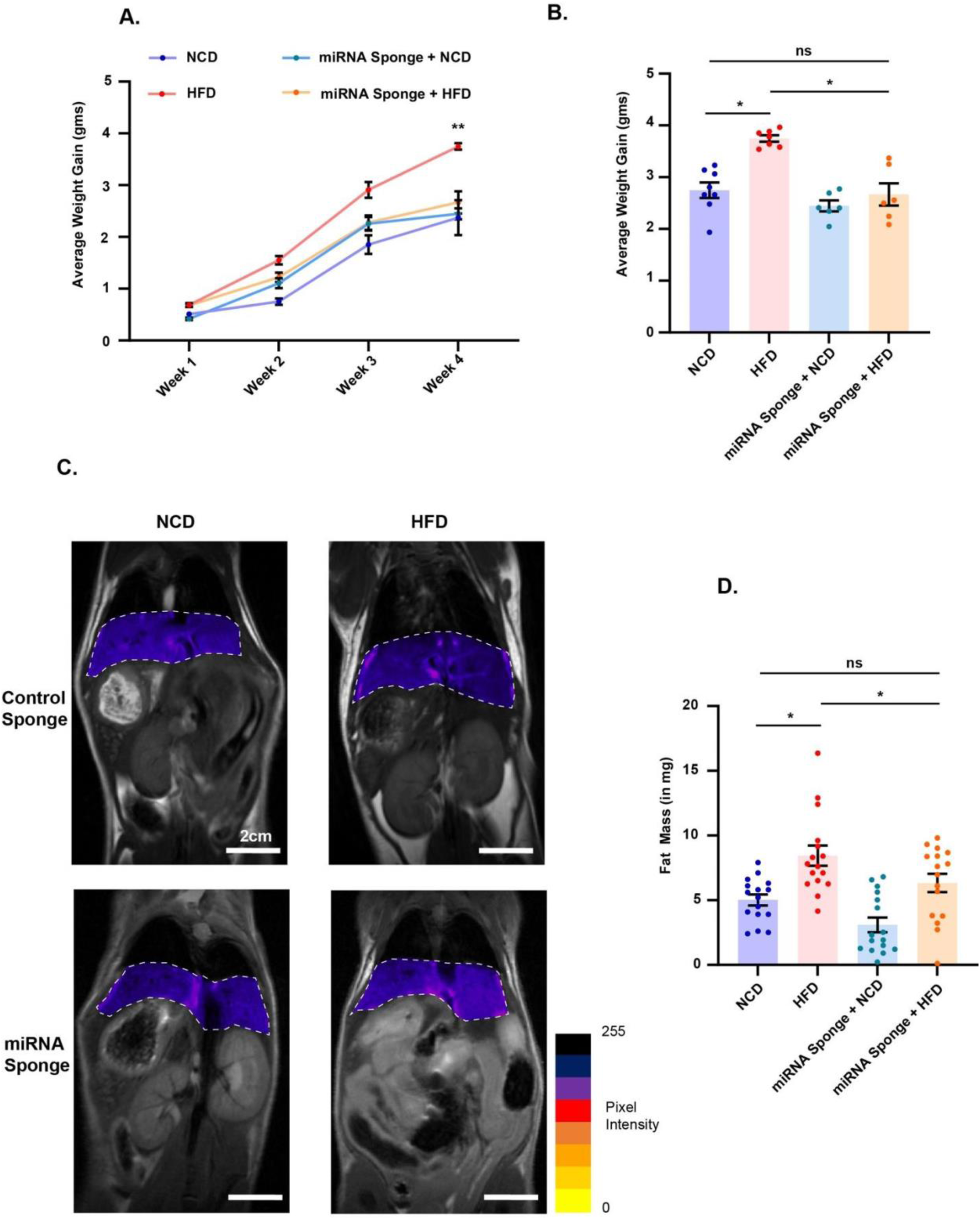
HFD -induced body weight gain and hepatic fat accumulation is prevented by inhibition of diet -regulated miRNAs. (A) Line plot of body weight gain measurements of mice fed with HFD or NCD, with or without the injections of sponge, for the duration of the feeding. N = 3 mice. **p < 0.0032. Data shown as mean ± SEM. Two tailed t-test with Welch’s correction. (B) Average body weight gain of mice fed with HFD or NCD, with or without the injections of sponge, at the end of the experiment (Weight at P75 – Weight at P45). N = 6-8 mice. *p < 0.0001. Data shown as mean ± SEM. Two tailed t-test with Welch’s correction. (C) Representative photomicrographs of T2-weighted MR Images of mice fed with HFD or NCD, with or without the injections of sponge, at P75. The highlighted hepatic region was selected for intensity analysis to determine fat mass in liver. (D) Quantification of hepatic fat mass analyzed from MR Images of mice fed with HFD or NCD, with or without the injections of sponge, at P75. n = 16 sections. N = 3 mice. *p < 0.0001. Data shown as mean ± SEM. Two tailed t-test with Welch’s correction.

## Discussion

Recent studies highlight how diet-induced neurogenesis and neuronal plasticity in the hypothalamus can profoundly influence feeding behaviour, energy expenditure, and systemic metabolism (4, 5, 37). Emerging evidence indicates that adult neurogenesis is dynamically regulated by both acute and chronic high-fat diet (HFD) exposure, contributing to divergent forms of obesity with distinct metabolic phenotypes. Notably, acute HFD feeding paradigms result in tanycytic neurogenesis from specific niches in the hypothalamus (5).

The current study attempts to reconcile physiological changes attributed to HFD exposure with HFD-induced tanycytic neurogenesis in the hypothalamus. Here, we elucidate the molecular underpinnings of neurogenesis from β2 tanycytes in the hypothalamus of young adult female mice following acute HFD feeding. We have identified a cohort of miRNAs that are expressed from different chromosomes, rather than a cluster of miRNAs from a single locus, after acute HFD feeding (Fig. 1). Network analysis revealed their shared mRNA targets, with GO terms implicating neurogenesis/differentiation regulators (Fig 2). A reporter assay confirmed the miRNA-mediated regulation of key hub transcripts, such as IGF1, CAMK2B, NTRK2 and ERBB4 (Fig. 3). HFD-induced miRNAs collectively favour the generation of functionally active (c-Fos+), orexigenic AgRP+ neurons over the anorexigenic POMC+ neurons (Fig. 4 - 6). Interestingly, simultaneous inhibition of all five HFD-induced miRNAs impaired the functional integration of newborn AgRP+ neurons into the hypothalamic feeding circuitry (Fig. 6 – 7); while also attenuating HFD-dependent body weight gain and hepatic fat accumulation (Fig. 8).

Several studies underscore the role of miRNA-mediated regulation in the modulation of hypothalamic feeding circuits with profound implications in obesity and metabolic control. Price *et al.* (35) found that miR-33 deletion in AgRP+ neurons increases their activity, boosting food intake and weight gain while causing metabolic dysregulation and worsening hepatic fat accumulation. LaPierre *et al.* (36) showed that deletion of miR-7 (a miRNA highly expressed in the hypothalamus) disrupts melanocortin signaling, promoting obesity. However, a holistic perspective on how diet-modified miRNAs concertedly function to induce neurogenesis and modify the feeding circuitry is unexplored (8, 38–41).

Our study resolves this void by identifying five HFD-induced miRNAs from β2 tanycytes in the MEm of adult hypothalamus. Network analysis revealed a unique set of mRNA transcripts that are combinatorially controlled by these five HFD-induced miRNAs; allowing us to inspect the regulatory pathways associated with neurogenesis, neuronal differentiation and diet-induced metabolic changes with greater resolution. These transcripts are positioned at key hubs within the network that can influence many nodes. GO analysis of the target mRNAs confirmed the existence of a neurogenic network that is regulated by HFD-induced miRNAs; with GO terms such as “response to hormone”, “neuron differentiation”, “regulation of cell migration” etc. showing maximum enrichment. Overall, a high degree of network modularity biased towards neurogenesis was observed in the target mRNAs of HFD-induced miRNAs, which was notably absent in the targets of NCD-induced miRNAs.

We developed an in-house miRNA sponge, which had complementary seed sequences against all five HFD-induced miRNAs aligned in tandem. The miRNA sponge had a greater degree of affinity to the miRNAs than their endogenous targets. Previous tanycyte ablation studies (5, 8) using X-ray irradiation lacked molecular specificity as they completely eliminated tanycytes. Our miRNA-sponge approach overcomes this limitation, revealing how specific miRNAs precisely regulate the translational landscape of the β2-tanycytic neurogenic niche at a molecular resolution. Notably, the development of a miRNA sponge serves as a proof-of-concept for RNA-based therapeutic strategies aimed at preventing obesity.

Modification of a neuronal circuitry may be achieved by the functional integration of newborn neurons into the pre-existing circuit; such neurons exhibit c-Fos expression (42). We found progenitors from β2 tanycytes differentiate into both orexigenic AgRP^+^ and anorexigenic POMC^+^ neurons after HFD feeding. However, inhibiting HFD-induced miRNAs significantly reduced AgRP+ neurons but didn’t affect POMC+ neuron generation, suggesting that HFD-induced miRNAs favour orexigenic neurogenesis. Expression of c-Fos in nascent AgRP^+^, but not POMC^+^ neurons suggest that these POMC^+^ neurons are not functionally active. A plausible reason being that the life of nascent POMC^+^ neurons under acute HFD-feeding paradigms are transient in nature. We postulate that β2 tanycyte-derived POMC^+^ neurons may act as a “brake” to HFD-induced metabolic changes. miRNAs upregulated under HFD favours the genesis and integration of orexigenic neurons into the feeding circuitry to create a sustained physiological demand for high fat containing diet. Acute HFD therefore fails to functionally integrate anorexigenic POMC neurons, favouring orexigenic-driven hyperphagia and weight gain, as mentioned in previous studies (5, 11). This impairment of functional POMC neuron integration is of potential therapeutic interest in countering the deleterious effects of high-fat diets.

This study offers two potential avenues of therapeutic intervention; a) targeting the miRNAs that are enriched by HFD using endogenously expressed miRNA sponges; b) overcoming molecular paradigms that inhibit the integration of nascent POMC+ neurons.

Although miRNA-dependent mechanisms of HFD-induced tanycytic neurogenesis has been characterized in young adult female mice, observations on further changes within tanycytic neurons in older animals is beyond the scope of this study. Future investigations on age-dependent and sex-dependent discrepancies in the molecular identities of HFD-induced newborn neurons may provide useful insights towards combating metabolic disorders such as obesity and hyperphagia.

Collectively, the findings from our study reveal the molecular underpinnings of an unexplored mechanism linking HFD-induced, miRNA-mediated neurogenesis in β2 tanycytes to weight gain and hepatic fat accumulation, that is driven by the selective integration of AgRP+ neurons into the feeding circuit.

## Materials and Methods

### Stereotaxic surgery for cannula implantation

Female C57BL6/j animals at P21 - P25 were used for the experiments. Cannulas were prepared from a 23G needle, modified for the appropriate ventral depth to be placed atop the skull as described in (43). They were fitted with a dummy copper wire inside to prevent the leak of CSF or the entry of external irritants into the lateral ventricle. This cannula structure was then sterilized before the surgeries.

Animals were anaesthetized with Ketamine (80mg per 1 kg of body weight) and Xylazine (20mg per 1kg of body weight) administered via subcutaneous injections. Post anaesthesia, the hairs were trimmed, and the animals were affixed onto the stereotactic instrument. The skull was exposed and the coordinates of the Bregma was determined. A hole for the cannula was drilled at the stereotaxic coordinates (Anterior +0.02cm; Lateral −0.08cm; Ventral −0.25cm w.r.t Bregma). Stereotaxic coordinates were obtained from the Mouse Coronal Atlas from the Allen Brain Atlas (mouse.brain-map.org). The cannula was lowered at the previously drilled site. Post the implant of the cannula, a small hole was drilled on the skull posterior to the cannula. A small screw was screwed onto the skull. Then, the skull was covered up with the dental cements and quick solidifying polymer. Then, the animals were allowed to recover from surgeries for 10-12 days, with regular changing of the sterile dummy copper wire inside the cannula. Animal experiments were performed with the approval of the Institutional Animal Ethics (IAEC) committee of National Brain Research Centre.

### Feeding and Intracerebroventricular/Intraperitoneal injections of BrdU

Animals were either started on a Rodent Diet with 60 kcal% fat diet (Research Diets, Inc. (RD #12492)) [HFD], or continued with the normal chow diet [NCD] at P45, and this was continued till they turned P75, when the animals were sacrificed for further experiments. Animals were given *ad libitum* access to the diet and water. All the animals used for miRNA expression analysis via TaqMan Arrays, were implanted with cannulas (n=3 animals for n=1 experiment for 3 experiments). These animals were injected with BrdU intracerebroventricularly at 2 injections a day of 10 µgms each from P45-P53. Mice used for immunostaining experiments were not implanted with cannulas (n=1 animal for n=3 experiments). These animals were injected with 10mg of BrdU per kg body weight intraperitoneally of the respective animals twice a day from P45-P53. Regular measurements of body weights were noted from P45-P75.

### Immunostaining

Animals were anaesthetized and perfused transcardially at P75 with 1X PBS / 4% Paraformaldehyde (PFA). The brains were then isolated and stored in PFA and processed with 30% Sucrose for cryosectioning. Brains were sectioned at 40 µm thickness and collected onto gelatin-coated slides for processing. Sections were then introduced to an HCl and Borate Buffer method for breaking the chromosomes and hybridized with anti-BrdU antibodies. They were also co-stained with anti-Vimentin, anti-Nestin, anti-POMC, anti-AgRP, anti-c-Fos and anti-NeuN antibodies as per the requirement of the different studies.

### Laser Capture Microscopy

Mice were anaesthetized and quickly perfused transcardially at P75 with 1X PBS on P75, and the brains were isolated, and flash frozen in liquid N_2_. Brains were sectioned at 12 µm thickness, and carried onto MembraneSlide 1.0 PEN (Zeiss). Sections were fixed with PFA, then stained for BrdU in RNase-free solutions, and conjugated with fluorescent tags to visualize under the microscope. Zeiss PALM MicroBeam Laser Microdissection microscope was used to visualize the fluorescent tags of BrdU. Manual ROIs were drawn around the fluorescent cells and they were micro dissected with the MicroBeam laser. The cells were collected onto miRNA lysis buffers and pooled for miRNA isolation using the Mirvana Total RNA isolation protocol (Applied Biosystems).

### TaqMan Low Density Array

The RNA from the captured neurogenic population was reverse transcribed using the Megaplex^TM^ RT Primers, Rodent Pools Set v3.0. Then, the cDNA was amplified using Megaplex^TM^ PreAmp Primers, Rodent Pool Set v3.0. The amplified cDNA was then used to analyze the miRNA expressions determined using TaqMan^TM^ Array Rodent MicroRNA A+B Cards Set v3.0. Data was then collected from 3 experiments, and average expression of the miRNAs were analyzed and plotted.

### miRNA target identification

Annotated and putative targets of diet responsive miRNAs were obtained from the databases of TargetScan (44), miRdb (45, 46) and PicTar (47). Common identified targets for each miRNA across all the databases were acquired. Then, the common targets for the cohort of all diet responsive miRNAs were also pooled into a single list of 1204 targets.

### Network Analysis

Interactions between the common targets of the HFD-enriched miRNAs were determined using the String database (https://string-db.org/). The interaction network of the targets was then analyzed in Matlab using Brain Connectivity Toolbox (24). All mRNA targets of the miRNAs were assigned as nodes of the network and the interactions between two nodes were considered as the edges of the network. Community analysis was performed using Louvain’s method to determine the community clusters of the nodes. Within module z-scores and participation coefficients were determined for each node and plotted. Nodes were determined to be hubs if their corresponding z-scores were > 2.

### Gene Ontology Analysis

Hubs selected by z-score analysis were used for the analysis of Gene Ontologies using PANTHER Database (http://pantherdb.org/) (doi:10.5281/zenodo.6399963). GO Enrichment for the hubs was obtained from the database. Fisher’s exact test was performed to test the significant enrichment of the transcripts in annotated datasets such as, GO: Biological Process and Panther Pathways, and the FDR values were generated using Panther statistical overrepresentation test. GO terms with significant fold enrichment were analyzed and plotted.

### miRNA sponge construct design

Bulged binding sites against the seed sequences of HFD enriched miRNAs (from miRBase (48)) were designed and synthesized (49) 6 sites for each of the 5 miRNAs (miRs −382, −145, −196, - 1894 and let-7b) were included in the synthesized construct (50). This construct was then incorporated into the ORF of the fluorescent marker tdTomato in Plasmid #59462 from Addgene.

An empty vector did not express the sponge sites. The plasmids were transfected into Neuro-2a cells for luciferase assays, and they were packaged into AAVs for injection into the hypothalamus for imaging. The designed sponge sequence is reported below with the bulged binding sites for miR-382, miR-145, miR-196, miR-1894 and let-7b in the sequences.

#### miRNA Sponge Fwd. (5’–3’)

AATTCAAAAAGTGCCACCGTGAATGATTATAAAAAGTGCCACCGTGAATGATTATAAAAAGTGCCACCGTGAATGATTATAAAAAGTGCCACCGTGAATGATTATAAAAAGTGCCACCGTGAATGATTATAAAAAGTGCCACCGTGAATGATTATCAAGAACATGCTTCCAGGAATTTATCAAGAACATGCTTCCAGGAATTTATCAAGAACATGCTTCCAGGAATTTATCAAGAACATGCTTCCAGGAATTTATCAAGAACATGCTTCCAGGAATTTATCAAGAACATGCTTCCAGGAATTTATATCAGGTGACCGATGTCGTTGTTATATCAGGTGACCGATGTCGTTGTTATATCAGGTGACCGATGTCGTTGTTATATCAGGTGACCGATGTCGTTGTTATATCAGGTGACCGATGTCGTTGTTATATCAGGTGACCGATGTCGTTGTTATCTCCCTTCGATTCTCCCTTGCTTATCTCCCTTCGATTCTCCCTTGCTTATCTCCCTTCGATTCTCCCTTGCTTATCTCCCTTCGATTCTCCCTTGCTTATCTCCCTTCGATTCTCCCTTGCTTATCTCCCTTCGATTCTCCCTTGCTTATGGGAAGGCGTGGGTTGTATAGTTATGGGAAGGCGTGGGTTGTATAGTTATGGGAAGGCGTGGGTTGTATAGTTATGGGAAGGCGTGGGTTGTATAGTTATGGGAAGGCGTGGGTTGTATAGTTATTTATGGGAAGGCGTGGGTTGTA TAGA

#### miRNA Sponge Rev. (5’–3’)

TCTATACAACCCACGCCTTCCCATAAATAACTATACAACCCACGCCTTCCCATAACTATACAACCCACGCCTTCCCATAACTATACAACCCACGCCTTCCCATAACTATACAACCCACGCCTTCCCATAACTATACAACCCACGCCTTCCCATAAGCAAGGGAGAATCGAAGGGAGATAAGCAAGGGAGAATCGAAGGGAGATAAGCAAGGGAGAATCGAAGGGAGATAAGCAAGGGAGAATCGAAGGGAGATAAGCAAGGGAGAATCGAAGGGAGATAAGCAAGGGAGAATCGAAGGGAGATAACAACGACATCGGTCACCTGATATAACAACGACATCGGTCACCTGATATAACAACGACATCGGTCACCTGATATAACAACGACATCGGTCACCTGATATAACAACGACATCGGTCACCTGATATAACAACGACATCGGTCACCTGATATAAATTCCTGGAAGCATGTTCTTGATAAATTCCTGGAAGCATGTTCTTGATAAATTCCTGGAAGCATGTTCTTGATAAATTCCTGGAAGCATGTTCTTGATAAATTCCTGGAAGCATGTTCTTGATAAATTCCTGGAAGCATGTTCTTGATAATCATTCACGGTGGCACTTTTTATAATCATTCACGGTGGCACTTTTTATAATCATTCACGGTGGCACTTTTTATAATCATTCACGGTGGCACTTTTTATAATCATTCACGGTGGCACTTTTTATAATCATTCACGGTGGCACTTTTTGAATT

### Luciferase reporter construct design

Primers to amplify the 3’UTR sequences of miRNA targets, IGF1, NTRK2, CAMK2B, ERBB4 and ANAPC11 were designed. The 3’UTRs were amplified from mouse brain RNA samples and were incorporated into the pMIR-Report vector into the 3’UTR of the luciferase reporter.

IGF1_F TTAAACAGTTAAGCTAGGAAGTGCAGGAAACAAGACC

IGF1_R ATCCTTTATTAAGCTCATCTTAGGCTCCAGGCTTTCG

NTRK2_F TTAAACAGTTAAGCTAGGCATCTCCCGTCTACCTG

NTRK2_R ATCCTTTATTAAGCTAGAGCGTCTCCTGTCTTTGTC

CAMK2B_F TTAAACAGTTAAGCTACGGCAAGTGGCAGAATGTAC

CAMK2B_R ATCCTTTATTAAGCTACCTGCATTGCCCAGAAGTC

ERBB4_F TTAAACAGTTAAGCTCGCCCTACAGACACCGGAATAC

ERBB4_R ATCCTTTATTAAGCTCCTTTCATGTAGACGTACCCAATCC

ANAPC11_F TTAAACAGTTAAGCTATGTGTCGCCAGGAGTGGAAG

ANAPC11_R ATCCTTTATTAAGCTAGTCTCCCAGAAGCACAAGTC

### Luciferase reporter assays

Neuro-2a cells were transduced with miRNA sponge, or control plasmid at 16 hrs post plating. Transfected cells were incubated in high glucose DMEM media (Gibco) with 5% fetal bovine serum and maintained at 5% CO_2_ and 37°C. Cells were simultaneously co-transfected with Gaussia luciferase containing the 3′UTRs of IGF1/ NTRK2/ CAMK2B/ ERBB4/ ANAPC11 and Firefly luciferase cloned in pMIR-Report plasmid as well as pRLTK. Luciferase activities were measured 48 hrs post transfection. Dual Luciferase Reporter Assay System (Promega) was used to measure the intensities of Gaussia and Firefly luciferases. Data was analyzed as the ratio of intensities from Gaussia luciferase to Firefly luciferase.

### AAV Preparation and Stereotaxic Injection into arcuate nucleus

Adenovirus was produced by cotransfection of 9.55 µgm of empty plasmid (expressing tdTomato by CAG promoter, #59462 from Addgene) or sponge plasmid (tdTomato with the sponge sequence inserted in the ORF under the CAG promoter), 9.55 µgm pAAV2/9n and 19.22 µgm pAdDeltaF6 in HEK293T cells. Calcium phosphate method of transfection was used to deliver the plasmids to HEK293T cells. Cells were incubated in low glucose DMEM media (Gibco) with 10% fetal bovine serum and maintained at 5% CO_2_ and 37°C. 72 hours after transfection, media was collected, and the cells were lysed and filtered with a 100-micron filter mesh. The media and lysed cells were pooled, and viral pellets were obtained using ultracentrifugation. Viral particles were resuspended in 1X PBS with 5% Glycerol.

Stereotaxic surgeries were performed on the mice aged P21-P25 to deliver the AAV containing miRNA sponge to the arcuate nucleus as mentioned above. Using a Hamilton syringe, 200nL of the AAVs were injected into the arcuate nucleus at the stereotaxic coordinates (Anterior −0.02 cm; Lateral −0.002 cm; Ventral −0.059 cm; w.r.t Bregma). After the surgery, animals were allowed to recover for 10-12 days before the feeding and injections were started.

### Confocal Imaging and Image Analysis

Brain sections stained and mounted were imaged using a Nikon A1 HD25 point scanning microscope with a Nikon 20X Objective at a 1024*1024 resolution of pixels. Optical sections of 1µm step sizes were captured for the entire brain sections. All captured images were of 16-bit pixel depth, and had uniform laser power, detector gain and pinhole diameter values.

TIFFs of raw images were selected from all channels expressing DAPI, BrdU, POMC, AgRP, c-Fos and NeuN. Using custom written MATLAB algorithms, all images were thresholded dynamically with their respective background noise. Thresholded images at the same optical section depths were used to quantify the underlying number of cells expressing in each channel as well as the number of colocalization between the channels. The data was then exported, pooled, analyzed and plotted for significance.

### miRNA Sponge Efficacy Analysis

Primary neuronal cultures were prepared from dissected mouse embryonic (E17) cortices as described previously (51). About 160 to 170 cells/mm^2^ were plated for verifying the efficacy of miRNA sponge expression. Neurons were grown *in vitro* in Neurobasal medium (Gibco) containing B27 supplements (Gibco) at 5% CO_2_/37°C. Adenoviruses expressing miRNA sponge or empty sponge vectors were transduced at DIV (days *in vitro*) 7. Neurons were fixed with 4% paraformaldehyde at DIV 21 and were counterstained with DAPI containing media.

Neurons were imaged using a Nikon A1 HD25 point scanning microscope with a Nikon 100X Objective at a 1024*1024 resolution of pixels. Optical sections of 0.2 µm step sizes were captured for the entire neuron. All captured images were of 16-bit pixel depth, and had uniform laser power, detector gain and pinhole diameter values.

Maximum Intensity Projections of each neuron were made as TIFFs from raw images and the average global pixel intensities of tdTomato fluorescence were computed using Matlab. The data was collected and pooled for statistical analysis.

### Magnetic Resonance Imaging and Analysis

All imaging procedures were conducted using a 7.0 T Bruker BioSpec 70/20 USR AV3HD small-animal MRI system (M/s Bruker BioSpin, Ettlingen, Germany), with Maximum gradient strength of 440 mT/m. A 70 mm quadrature volume coil was employed for both radiofrequency transmission and reception.

#### Preparation

Mice at P75 were anaesthetized with Ketamine (80mg per 1 kg of body weight) and Xylazine (20mg per 1kg of body weight) administered via subcutaneous injections. The animals were positioned in a prone orientation (head-first in the scanner) with the abdomen centered within the coil. Core body temperature was maintained at 36–37°C utilizing a warm-air feedback system, and respiration was monitored via a pneumatic pillow sensor. Prospective respiratory gating was applied to all sequences, with acquisitions triggered during end-expiration.

#### Imaging protocol

A respiratory-gated, two-dimensional T₂-weighted TurboRARE sequence (Bruker: RARE) was utilized, with repetition times (TR) ranging from 1411-1852ms and an echo time (TE) of 27ms. The echo train length (RARE factor) was conFig.d to 8, with a flip angle of 90°. Images were acquired using a 320 × 320 reconstructed matrix, a 100% phase field of view, and an in-plane resolution of approximately 0.10–0.125 mm, adjusted according to the field of view for each subject. Both slice thickness and interslice spacing were set to 1.0 mm, with 8 signal averages (NEX) and a pixel bandwidth of 244 Hz/pixel. The phase-encoding direction was designated as either ROW or COL, contingent on anatomical geometry, to minimize motion ghosting. All acquisitions employed prospective respiratory gating, triggered at end-expiration, and were conducted in ParaVision Acquisition 6.0.1. For fat-suppressed acquisitions, the same sequence geometry and parameters were maintained, with the addition of spectral fat-suppression applied at the lipid resonance frequency prior to excitation to selectively attenuate the fat signal. Each gated acquisition lasted approximately 3–6 minutes, depending on the respiratory rate.

#### Imaging Analysis

Raw Bruker data were converted to NIfTI format using dcm2niix (v1.0.20240202), ensuring complete preservation of orientation metadata. Intensity histograms were calculated from NIFTI format images. Using custom program in Matlab, Otsu’s threshold was applied to the images and voxels with high intensities (> Image Threshold) were considered as “fat voxels” and intermediate intensities were considered as “lean voxels”. Liver masks were generated by manual ROIs to isolate hepatic fat from visceral fat. Voxel volumes were determined, and fat mass was calculated with known density for fat tissue (0.9 gm/cm^3^).

### Statistics

Statistical analysis was performed for all experiments. miRNA differential expressions were analyzed for significance using two-tailed students’ t-test and post hoc determination of false discovery rates using the Benjamini-Hochberg method (52). Luciferase reporter assays and intensity analysis for miRNA sponge expression were analyzed for significance using unpaired two-tailed t-test with Welch’s correction. Unpaired t-test with Welch’s correction was used to determine the significance for all immunostaining experiments including determination of orexigenic and anorexigenic fate and their functional integration in the hypothalamic feeding neural circuit. The functional activity of neurogenic cells was also analysed by unpaired t-test with Welch’s correction.

## Acknowledgments

We thank Addgene for AAV vectors. We thank Ananya Ghosh and Santhosh Kumar for the technical assistance. This study was funded by the Department of Biotechnology, Government of India (BT/PR8793/AGR/36/749/2013) and NBRC core fund to Sourav Banerjee.

## Author Contributions

<u>Conceptualization:</u> Sourav Banerjee

<u>Data curation:</u> Balakumar Srinivasan, Sarbani Samaddar, Himanshu Singh

<u>Data analysis:</u> Balakumar Srinivasan, Sarbani Samaddar, Himanshu Singh

<u>Writing – original draft:</u> Balakumar Srinivasan, Sarbani Samaddar, Sourav Banerjee

<u>Writing – review & editing:</u> Sarbani Samaddar, Sourav Banerjee

<u>Funding acquisition:</u> Sourav Banerjee.

## Data Availability

All data and codes used for analysis are available at https://figshare.com/s/abef061d9d61a06f3dd7

## Supporting Information

**Fig S1:**
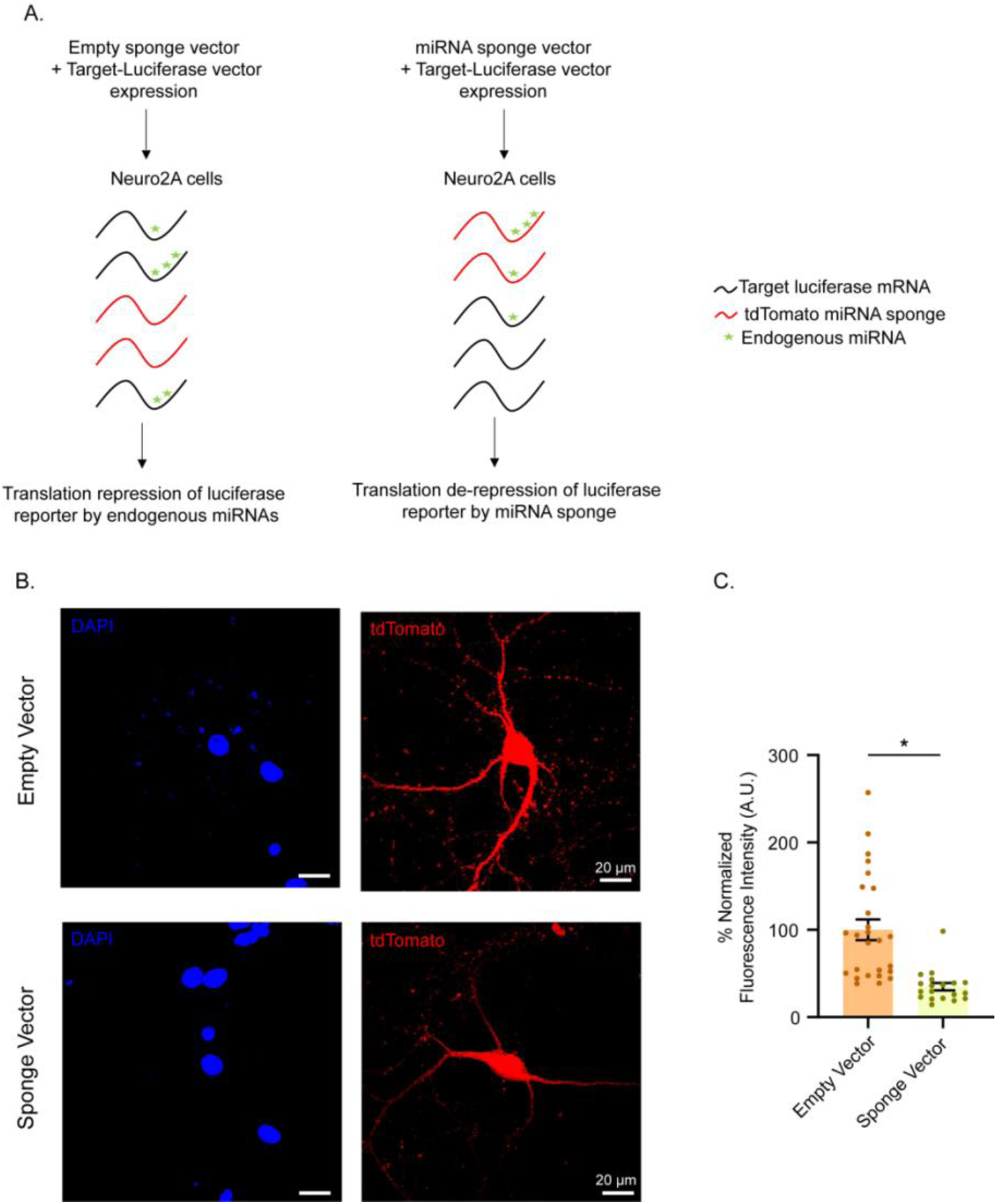
Inhibition of diet -regulated miRNAs by sponge. (A) Schematic for miRNA sponge-mediated inhibition of miRNA-mRNA target interaction as measured by reporter assay. (B). Photomicrograph representing miRNA sponge activity in primary cortical neurons following AAV-mediated miRNA sponge expression. Scale as indicated. (C) Quantitation of inhibition of tdTomato expression by miRNA sponge in primary cortical neuron. N = 19 – 26 neurons from 3 independent experiments. *p<0.0001. Two-tailed t-test with Welch’s correction. Data shown as mean ± SEM.

